# Enzymatic Assembly for CRISPR Split-Cas9 System: The Emergence of a Sortase-based Split-Cas9 Technology

**DOI:** 10.1101/2024.02.29.582732

**Authors:** Seyed Hossein Helalat, Helga Thora Kristinsdóttir, Astrid Dolinger Petersen, Rodrigo Coronel Téllez, Mads Nordlund Boye, Yi Sun

## Abstract

CRISPR-Cas9 has been widely used in scientific research and medical investigations as a pioneering technology. However, challenges such as the large size of the Cas9 sequence and potential off-target effects have impeded its widespread adoption. In response, various alternatives, such as split-Cas9 technology, have emerged. Split-Cas9 systems allow the large Cas9 sequence to be divided into two segments to aid in the delivery of the enzyme. Nevertheless, challenges persist in achieving precise control over the timing and location of Cas9 reassembly and activity to minimize off-target effects. This study presents an enzymatic-based split-Cas9 system, introducing a new approach utilizing the Sortase enzyme for the reconstitution of the full Cas9 protein. The developed method eliminates the need for chemical or physical induction and allows for precise genome editing in specific cells through the utilization of various specific promoters or targeted drug delivery. Experimental validation of the enzymatic system was conducted in *E. coli*, HEK cells, and Jurkat cells, demonstrating successful assembly and activity of the assembled Cas9 enzyme. In addition, this study explored the incorporation of nuclear localization signals, the evaluation of inducible promoters, and the delivery of the system’s components in mRNA or protein form. Furthermore, we investigated the potential of S/MAR minicircle technology instead of viral vectors within the system. Overall, we highlighted the feasibility and utility of the Sortase-based split-Cas9 system to enhance control and efficiency compared to traditional CRISPR-Cas9 approaches. Additionally, this study revealed the potential of using the Sortase enzyme for posttranslational modifications and protein assembly in human cells.

## Introduction

CRISPR-Cas9 technology, a renowned user-friendly system, has significantly expanded the horizons of scientific research and medical interventions.^1^ Its versatile applications include high-throughput screening, genomics research, and advancements in diverse medical fields, including the treatment of infections, genetic disorders, and cancers, as well as the manipulation of immune cells and contributions to regenerative medicine.^2–4^ Despite its remarkable success, the CRISPR-Cas9 system is not without recognized drawbacks.^5,6^

A significant hurdle arises from the considerable size of the CRISPR-Cas9 sequence, exceeding 4 kb, which presents challenges for its delivery with commonly used adenoviral vectors.^7^ Additionally, occasional off-target effects associated with the system may result in undesired gene mutations. Furthermore, nonspecific gene regulation *in vivo* in untargeted cells represents another challenge. This finding emphasizes the importance of precise control over the CRISPR-Cas9 system to mitigate unintended consequences and optimize its therapeutic potential.^6,8^

In response to these challenges, various strategies have been developed, such as employing drug delivery methods involving nanoparticles for Cas9 enzyme (or RNP) delivery,^9^ creating inducible or specific Cas9 expression systems,^10^ and devising split-Cas9 systems.^7^ Among these approaches, split-Cas9 has emerged as a particularly promising method. This technique effectively diminishes cargo size by dividing the protein, whereby the two Cas9 fragments are separately introduced and expressed, rendering it inactive until reconstitution. Several methodologies have been developed in this area, including incorporating light- or chemically-inducible protein domain linkers and leveraging an intein-based assembly system.^11–13^

With respect to inducible systems, which act as a switch for Cas9 protein assembly or gene expression, various chemically and light-inducible CRISPR-Cas9 systems have been developed. However, applying these systems *in vivo* poses challenges such as potential cytotoxicity of chemicals, limited spatial resolution, the duration of chemicals remaining in the body, poor penetration of light into deep tissues, relatively low induction efficiency, and the requirement for manual control of the systems.^1,13,14^ In intein-based systems, inteins are integrated into split-Cas9, promoting the assembly of split parts to form an active Cas9. Although inteins may alleviate constraints linked to viral delivery,^15^ they do not fully resolve the issue of off-target effects. As such, there is a demand for new split-Cas9 technologies as alternatives.

Moreover, due to the drawbacks associated with viral vectors, including limited cargo capacity, immune responses, manufacturing complexity, risk of insertional mutagenesis, and potential for off-target effects, alternative methods for gene expression in mammalian cells have emerged.^16,17^ One such method is the use of minicircle (MC) plasmids for gene therapy, which enables long-term transgene episomal expression both *in vitro* and *in vivo*. This advantage stems from the absence of bacterial sequences, typically characterized by CpG dinucleotides, which are known triggers for vector-silencing mechanisms.^18^ Additionally, incorporating scaffold/matrix attachment regions (S/MARs) into these plasmids facilitates the binding of episomal vectors to chromosomal scaffolds during cell division, ensuring persistent expression and stability across numerous cellular generations.^19,20^ Given the promising features of MC plasmids, exploring their potential for split-Cas9 expression would also be highly beneficial.

In this work, we developed an innovative sortase-based split-Cas9 system utilizing an enzymatic assembly approach. Sortase A (SrtA), an epitope-specific enzyme, is a small transpeptidase derived from *Staphylococcus aureus* that serves as a powerful tool for the precise ligation of peptides and proteins. In this catalytic process, SrtA cleaves the specific amino acid recognition motif LPXTG at the threonine residue. Subsequently, upon the release of the glycine residue and all downstream amino acids, a sortase-bound thioester is formed and concurrently ligated to the nucleophile residue of the other protein, typically represented by the α-amino group of the N-terminal poly-glycine.^21,22^ Within this system, sortase-mediated trans-splicing occurs solely upon co-expression or delivery of SrtA, leading to the reconstitution of the full Cas9 protein (Figure 1).

**Figure 1.**
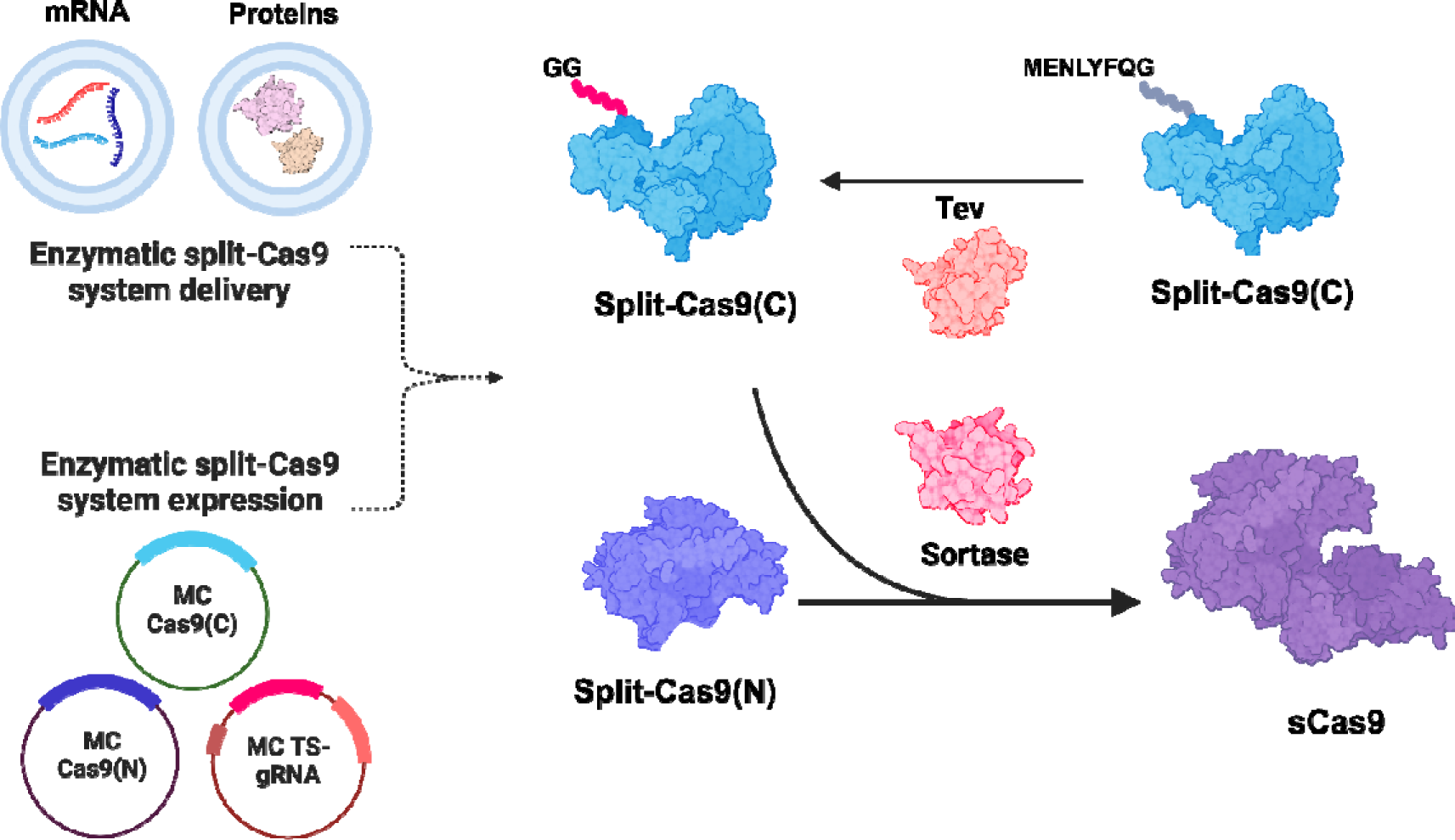
Enzymatic assembly of the split-Cas9 system. In this approach, the N-terminal region of the Split-Cas9(C) protein is cleaved by the Tev protease to generate a poly-G sequence at the N-terminus. Subsequently, the sortase enzyme ligates the LPXTG peptide of Split-Cas9(N) to the poly-G sequence of Split-Cas9(C), resulting in the generation of active sCas9. Additionally, various components of this system can be delivered via nanoparticles or expressed using episomal or viral vectors.

This system addresses the challenges associated with the intracellular delivery of Cas9 cargo and has the potential to reduce off-target effects associated with Cas9 activity in unintended cells. It offers customizable genome editing, which is particularly crucial when targeting a specific cell requires involving more than two activator input parameters, and represents a more controllable CRISPR-Cas9 system facilitated by an efficient and precise assembly mechanism. Additionally, the linker peptide for SrtA is a short peptide with minimal impact on Cas9 activity.

We also explored various strategies within the split-Cas9 system to showcase its potential and enhance control over Cas9 activity, particularly when utilizing MC plasmids. Our approach involved testing the expression of split components and assembly enzymes using two- and three-vector models, each controlled by distinct promoters. Additionally, we investigated the combination of mRNA delivery alongside simultaneous expression of other elements within the system. Furthermore, we assessed the feasibility of delivering assembly enzymes in protein form to cells expressing other system components. This investigation aimed to evaluate the potential utility of this system for precisely targeting specific cells in genome editing applications.

## Materials and Methods

### Strains and plasmid preparation

For cloning and plasmid amplification, *E. coli* DH5α was utilized. Prior to transformation, the cells were incubated overnight at 37°C and 250 rpm and then rendered chemically competent with CaCl_2_. Single colonies were selected and cultured in LB medium supplemented with either 50 µg/mL kanamycin or 100 µg/mL ampicillin at 37°C and 250 rpm.

To evaluate the performance of the split-Cas9 assembly system, we initially assessed its functionality in *E. coli* using the pET-srt Cas9 and pET-srt split573 plasmids. These plasmids were constructed by cloning genes encoding sortase, Tev protease, and Cas9/split-Cas9 into the pETDuet-1 vector backbone. Additionally, for protein delivery purposes, we constructed the pET-srt-histag plasmid to express the SrtA protein in *E. coli*.

Similarly, to assess the system in mammalian cells, codon-optimized genes were inserted into S/MAR minicircle plasmids. To generate the MC Cas9(C), MC Cas9(N), and MC TS-gRNA plasmids, distinct genes were cloned and inserted into S/MAR minicircle plasmids. Modifications, such as exchanging gRNAs in plasmids and adding a nuclear localization signal (NLS) to split-Cas9, sortase, and Tev protease, were achieved by amplifying and linearizing the plasmid using primers with corresponding sequences and overhangs, employing the GeneArt™ Gibson Assembly HiFi Master Mix (Thermo Fischer Scientific). Phusion Hot Start II High-Fidelity PCR Master Mix (Thermo Fischer Scientific) was used for PCR reactions.

For plasmid extraction, the NucleoSpin Plasmid Mini kit (MACHEREY-NAGEL) was used for minipreparations, while the QIAGEN Plasmid Plus Maxi kit was used for maxipreparations. The DNA concentration was determined using a NanoDrop™ 2000/2000c spectrophotometer (Thermo Scientific™). To confirm the DNA sequence of all plasmid constructs, samples were prepared accordingly and sent to Eurofins Scientific for sequencing utilizing the Mix2Seq Kit NightXpress.

### Protein modeling and *in silico* analysis

To determine the optimal sites for splitting the Cas9 enzyme, we identified three suitable locations positioned near the midpoint of the protein to create smaller split-form expression cassettes. Specifically, we targeted amino acids 297, 573, and 968 for the split sites. Models of the split proteins were then generated using the SWISS-MODEL, AlphaFold2, and I-Tasser servers.^23,24^ Each top rank (Rank 1) PDB structure was cleaned with the ‘RepairPDB’ command to prepare the PDB files for optimal FoldX analysis. Subsequently, the stability of these split proteins was evaluated using FoldX 5.0 software.^25,26^ The total energy of each construct’s score function was calculated and normalized by the length of each construct to obtain the free energy per residue, providing a basis for comparing the stability of the constructs.

Moreover, considering that the LPXTG and poly-G peptides are short, inactive peptides with no inherent affinity, it was assumed that these peptides should be easily accessible. The accessibility of the N-terminal and C-terminal sites of the split proteins was assessed using the generated models and the NetSurfP-2.0 tool.^27^ Additionally, visualization of the structures and alignment of the models were performed using PyMol software version 2.5.2.

### S/MAR Minicircle plasmid preparation

Minicircles were generated from the full-sized parental minicircle using PhiC31 integrase, which facilitated recombination between the PhiC31 attB and attP sites on the parental plasmid. This process yielded two products, the minicircle, which lacked any bacterial backbone DNA sequences, and the remaining part of the parental plasmid. To eliminate the second plasmid with a bacterial backbone in *E. coli*, the I-SceI endonuclease was employed to recognize and degrade the I-SceI sites in this plasmid, which led to its degradation. For efficient minicircle production, the *E. coli* ZYCY10P3S2T strain was utilized. This strain harbors an arabinose-inducible system for simultaneous expression of the PhiC31 integrase and the I-SceI endonuclease. Additionally, the ZYCY10P3S2T strain carries a robust arabinose transporter, the LacY A177C gene. Therefore, by supplementing the media with arabinose and inducing the expression of the PhiC31 integrase and endonuclease genes, the parental minicircle plasmid was separated into individual minicircles and plasmids harboring the bacterial backbone.

To execute the described procedure and generate minicircle plasmids for this study, we utilized the MC-Easy™ Minicircle DNA Production Kit (System Bioscience, SBI). Briefly, we initiated the process by inoculating 2 mL of LB-Kana medium with the ZYCY10P3S2T *E. coli* strain previously transformed with the minicircle plasmid, followed by incubation for 1 hour at 30°C and 250 rpm. Subsequently, 2 mL of culture was used to inoculate 200 mL of LB-Kana medium, which was then incubated for 16 hours at 30°C and 250 rpm. After the growth period, 200 mL of induction medium containing arabinose was added, and the culture was shaken for an additional 3 hours at 30°C. Then, the temperature was increased to 37°C for an additional 1 h, after which the cells were harvested for mini/maxiprep plasmid extraction.

To verify the removal of the bacterial backbone sequence, we examined the size and pattern of the purified plasmids through restriction enzyme digestion and analyzed the samples by agarose gel electrophoresis.

### Evaluation of the system in *E. coli* and protein purification

For the expression of the split-Cas9 and Cas9 plasmids, the *E. coli* BL21 strain was employed. Following transformation, the cells were cultured in LB medium supplemented with 100 µg/mL ampicillin and incubated at 37°C with agitation at 250 rpm until they reached an approximate OD600 of 0.8. Subsequently, induction was carried out by adding 0.5 mM IPTG, and the culture was left overnight at 30°C with agitation at 250 rpm.

For protein purification, as both normal Cas9 and assembled split-Cas9 contained His-tags, the proteins were initially purified using nickel beads. After centrifugation of the IPTG-induced cultures at 4500 × g for 15 minutes and removal of the supernatant, the cell pellet was lysed by resuspension in B-PER buffer (Thermo Fischer Scientific). Subsequently, HisPur™ Ni-NTA Resin (Thermo Fischer Scientific) was prepared and utilized for protein purification according to the manufacturer’s instructions. To enhance purity, size exclusion chromatography was employed using a Superose 6 (10/300) GL column (Cytiva) in PBS buffer (0.5 mL/min) on an ÄKTA Pure machine.

The assembly of split-Cas9 by our system was analyzed based on molecular weight using SDS[PAGE to assess the presence of the Cas9 protein at the expected size (≈150 kDa). For this purpose, the samples were prepared by adding Laemmli SDS sample buffer (4X, Thermo Fischer Scientific) and heating at 95°C for 5 minutes. Protein separation was performed using 12% Mini-PROTEAN® TGX Protein Gels (Bio-Rad) alongside PageRuler™ Prestained Protein Ladder (Thermo Fischer Scientific). Subsequently, the gels were stained with Bio-Safe™ Coomassie Stain (Bio-Rad) for 1 hour on an orbital shaker, followed by destaining with distilled water.

To investigate the activity of the assembled split-Cas9 enzyme in *E. coli*, the enzyme was purified and tested in Hek-Luc cells. Additionally, for the protein delivery assessment, we expressed and purified the SrtA-His tag enzyme using the same method. For the Tev protease protein required for this step, AcTEV™ Protease (Thermo Fischer Scientific) was used. For buffer exchange or protein concentration purposes, we employed Amicon® Ultra15 Centrifugal Filter Units (Merck) with 15 and 50 kDa cutoffs.

### Cell culture and gene delivery to mammalian cells

Normal HEK293T (Hek) and HEK293-Luc (BioCat GmbH, SL039-GVO-GC) cells were cultured at 37°C with 5% CO_2_ in DMEM supplemented with 10% fetal bovine serum (FBS) and 1% PenStrep (100 U/mL penicillin and 100 μg/mL streptomycin). Jurkat cells were cultured under the same conditions but in RPMI medium. Prior to the experiments, all the cell cultures were tested for mycoplasma contamination.

For delivering plasmids to HEK cells, both lipofection and electroporation techniques were employed. Jurkat cells were exclusively transfected via electroporation.

All DNA plasmids were transfected into Hek-Luc cells using Lipofectamine™ LTX Reagent (Thermo Fisher Scientific). Cells were seeded 24 hours prior to transfection to ensure high viability and suitable confluency. Subsequently, the DNA-lipid complex was added to the cells and incubated at 37°C with 5% CO_2_ for 3-6 days.

For electroporation of Jurkat cells, the Lonza™ P3 Primary Cell 96-well Nucleofector™ Kit (Lonza) was utilized. Briefly, after the cells were washed with PBS buffer, they were suspended in P3 Primary Cell Nucleofector™ Solution (Lonza) and mixed with the desired amount of plasmid per reaction (ranging from 0.2 to 5 μg). This cell-plasmid mixture was then loaded into the electroporation wells. Electroporation was carried out using the Jurkat CL120 program on a 4D-Nucleofector® Core Unit (Lonza) machine. Following electroporation, additional media was added to the cells, which were subsequently incubated at 37°C with 5% CO_2_ for 3-5 days.

Notably, in this study, 150,000 cells were used for each reaction for both cell lines. Different plasmid ratios and concentrations (ranging from 0.5 µg to 2.5 µg) were tested to evaluate the effect of varying plasmid concentrations on knockout efficiency.

Jurkat cell activation was induced using Gibco™ Dynabeads™, Human T-Activator CD3/CD28 for T-cell expansion and activation.

### Protein delivery

To simulate the delivery of SrtA and Tev proteins to cells using nanoparticles, we used Pierce™ Protein Transfection Reagent (Thermo Fisher Scientific) to generate nanoparticles with efficient transfer capabilities for these proteins. This product facilitates the creation of cationic lipid-based carriers encapsulating SrtA and Tev enzymes, enabling their entry into cells either by direct fusion with the plasma membrane or through endocytosis, followed by fusion with the endosome and subsequent release of the encapsulated proteins into the cytoplasm. The carriers were prepared following the manufacturer’s protocol, with 100 ng of protein utilized for each 10 µl of Pierce reagent during the hydration step.

### Preparation of gRNAs and RNP complexes

Preparation of double-stranded gRNAs was carried out to establish positive controls using a commercial Alt-R® Cas9 (Integrated DNA Technologies) and to test purified split-Cas9 expressed and assembled in *E. coli*. Itgal-crRNA, luc-crRNA, and tracrRNA were procured from Integrated DNA Technologies.

The crRNA and tracrRNA were diluted with nuclease-free IDT buffer to achieve a concentration of 200 μM. Subsequently, they were mixed in equimolar concentrations and heated at 95°C for 5 minutes to facilitate hybridization. For the formation of ribonucleoprotein (RNP) complexes, two protocols were employed, both demonstrating similar efficacy and specificity. In the first protocol, a total volume of 5 μL was prepared by combining 2.1 μL of PBS, 1.2 μL of gRNA, and 1.7 μL of Alt-R® Cas9. Conversely, the second protocol involved the preparation of a 2.2 μL mixture consisting of 1.2 μL of gRNA and 1 μL of Alt-R Cas9 or our expressed Cas9/split-Cas9. After assembling the RNP complexes according to the prescribed protocols, the mixtures were incubated at room temperature for 15 minutes. The final mixtures were then added to the cells for the electroporation procedure.

To knock out the luciferase gene in Hek-Luc cells, two variations of gRNAs targeting *luc* were designed and tested: luc1 (GGCTATGAAGAGATACGCCC) and luc2 (GGATCTACTGGGTTACCTAA). For the knockout of the *itgal* gene in Jurkat cells, the gRNAs Itg1 (GATGATGATAAGCACTTTGG) and Itg2 (GGCTACCCACTCGGGCGGTT) were designed and tested.

### Western blot

Hek cells transfected with the respective split-Cas9 plasmids were lysed 24-72 hours post-transfection by adding Laemmli buffer containing 10 mM DTT, followed by either boiling on a heating block or sonication. The protein extract samples were then loaded onto an SDS-PAGE gel, which was subsequently transferred to a polyvinylidene fluoride (PVDF) membrane. To activate the PVDF membrane, it was first treated with 96% ethanol and then placed in deionized water. The transfer sandwich, comprising the PVDF membrane and the SDS-PAGE gel, was assembled and transferred to a chamber filled with transfer buffer (20% absolute ethanol, 12.4 mM Tris, and 96 mM glycine). The transfer process was conducted at 140 mA for 90 minutes. Subsequently, the membrane was blocked with blocking buffer (5.73 mM skim milk in 50/50 PBS-Tween buffer) for 1 hour on an orbital shaker. After blocking, the membrane was washed three times with PBS-Tween buffer for 5 minutes per wash with shaking. The washed membrane was then incubated overnight at 5°C with a 6x-His-Tag monoclonal antibody (Invitrogen) diluted 2500-fold in PBS-Tween. A His-tag antibody solution was used because the proteins carry a 6xHis-tag, facilitating antibody binding. Following incubation, the membrane was washed three times with PBS-Tween before being subjected to a chemiluminescence (CL) reaction. For the CL reaction, 500 µL of each reagent from Pierce™ ECL Western Blotting Substrate (Thermo Fisher Scientific) was mixed, and the substrate was added to the membrane. Finally, the membrane was transferred to a CL detector for imaging.

### Flow cytometry analysis

The cells were washed with PBS (Gibco) and resuspended in PBS. Cell viability staining was performed using Near-IR Live/Dead (NIR, Life Technologies, L10119) kit. Surface staining was carried out using a fluorochrome-labeled PE-conjugated anti-human CD11a (HI111) monoclonal antibody obtained from BioLegend.

The stained samples were analyzed on a Fortessa LSR flow cytometer (BD), and the data were analyzed using FlowJo V.10.9.0 software. Statistical significance was determined with a two-sided P value <0.05 considered statistically significant.

### Luciferase activity assay

Luciferase emissions from transfected Hek-Luc cells were measured using the Pierce® Firefly Luciferase Glow Assay Kit (Thermo Fisher Scientific). To initiate cell lysis, the media was aspirated from the cells, followed by rinsing with PBS. Subsequently, the cells were mixed with 1X Cell Lysis Buffer on a platform shaker for 15 minutes. The resulting cell lysates were then transferred to a white opaque 96-well plate, and D-Luciferin solution with Pierce™ Firefly Signal Enhancer (100×) was added to each well.

Bioluminescence emitted upon luciferase binding to its substrate luciferin to form oxyluciferin was measured using a microtiter plate reader. The instrument quantified bioluminescence in relative light units (RLUs).

**Table 1.** Constructed plasmids.

| Name | Harboring genes | Source |
| --- | --- | --- |
| Parental Minicircle-MN501A1 | *CMV-MCS-EF1α-GFP, Kana resistance* | SBI |
| MC plasmid | *CMV-MCS, Kana resistance, S/MAR* | This study |
| pETDuet-1 | *Inducible T7 promoter, Amp resistance* | Novagen |
| pET-srt Cas9 | *SrtA, TEV, Cas9* | This study |
| pET-srt split573 | *SrtA, TEV, C-ter Cas9, N-ter Cas9* | This study |
| **pET-srt-histag** | *SrtA-His tag* | This study |
| **MC TS-luc** | *SrtA, TEV, mCherry, luc-gRNA* | This study |
| **MC TS-luc-srt-NLS** | *SortA-NLS, TEV, mCherry, luc-gRNA* | This study |
| **MC TS-luc-TEV-NLS** | *SortA, TEV-NLS, mCherry, luc-gRNA* | This study |
| **MC TS-luc-TEV-NLS** | *SortA-NLS, TEV-NLS, mCherry, luc-gRNA* | This study |
| **MC TS-itgal** | *SortA, TEV, mCherry, itgal-gRNA* | This study |
| **MC TS-itgal-srt-NLS** | *SortA-NLS, TEV-NLS, mCherry, itgal-gRNA* | This study |
| **MC TS-itgal-TEV-NLS** | *SortA, TEV-NLS, mCherry, itgal-gRNA* | This study |
| **MC TS-itgal-srt and TEV-NLS** | *SortA-NLS, TEV-NLS, mCherry, itgal-gRNA* | This study |
| **MC NFAT-TS- itg** | *NFAT promoter, SrtA, TEV, mCherry, itg-gRNA* | This study |
| **MC Cas9-luc** | *Normal Cas9, luc-gRNA* | This study |
| **MC Cas9-itg** | *Normal Cas9, itgal-gRNA* | This study |
| **MC Cas9(C)** | *Cas9(C)-NLS* | This study |
| **MC Cas9(C)-GFP** | *Cas9(C)-NLS, GFP* | This study |
| **MC Cas9(N)** | *Cas9(N)* | This study |
| **MC Cas9(N)-GFP** | *Cas9(N), GFP* | This study |
| **MC Cas9(N)-NLS** | *Cas9(N)-NLS* | This study |
| **MC Cas9(N)-TS-itg** | *Cas9(N), SortA, TEV, mCherry, itgal-gRNA* | This study |
| **MC Cas9(N)-TS-luc** | *Cas9(N), SortA, TEV, mCherry, luc-gRNA* | This study |

## Results

### Split-Cas9 variants

To identify the most suitable split variant for our system, we selected three distinct cleavage sites within the Cas9 protein at positions 297, 573, and 968. As no pre-existing PDB structures were available, *de novo* PDB structures of each Cas9 split variant were created and investigated *in silico*. Our analysis focused on evaluating the stability of each Cas9 split variant, revealing that there was no dramatic difference among the different split-Cas9 proteins. However, position 573 exhibited better stability on average, accommodating both the N-terminal Cas9(N) and Cas9(C) fragments (Figure S1A). Additionally, considering that the LPXTG and poly-G peptides are short, inactive peptides lacking inherent affinity, it was hypothesized that these peptides would be readily accessible. The accessibility of both the N-terminal and C-terminal sites of the split proteins was assessed, with *in silico* results indicating that position 573 demonstrated superior accessibility. Moreover, this position is closer to the midpoint of the Cas9 sequence, which could streamline the delivery of shorter gene segments through various systems (Figures 2A, and S1B).

**Figure 2.**
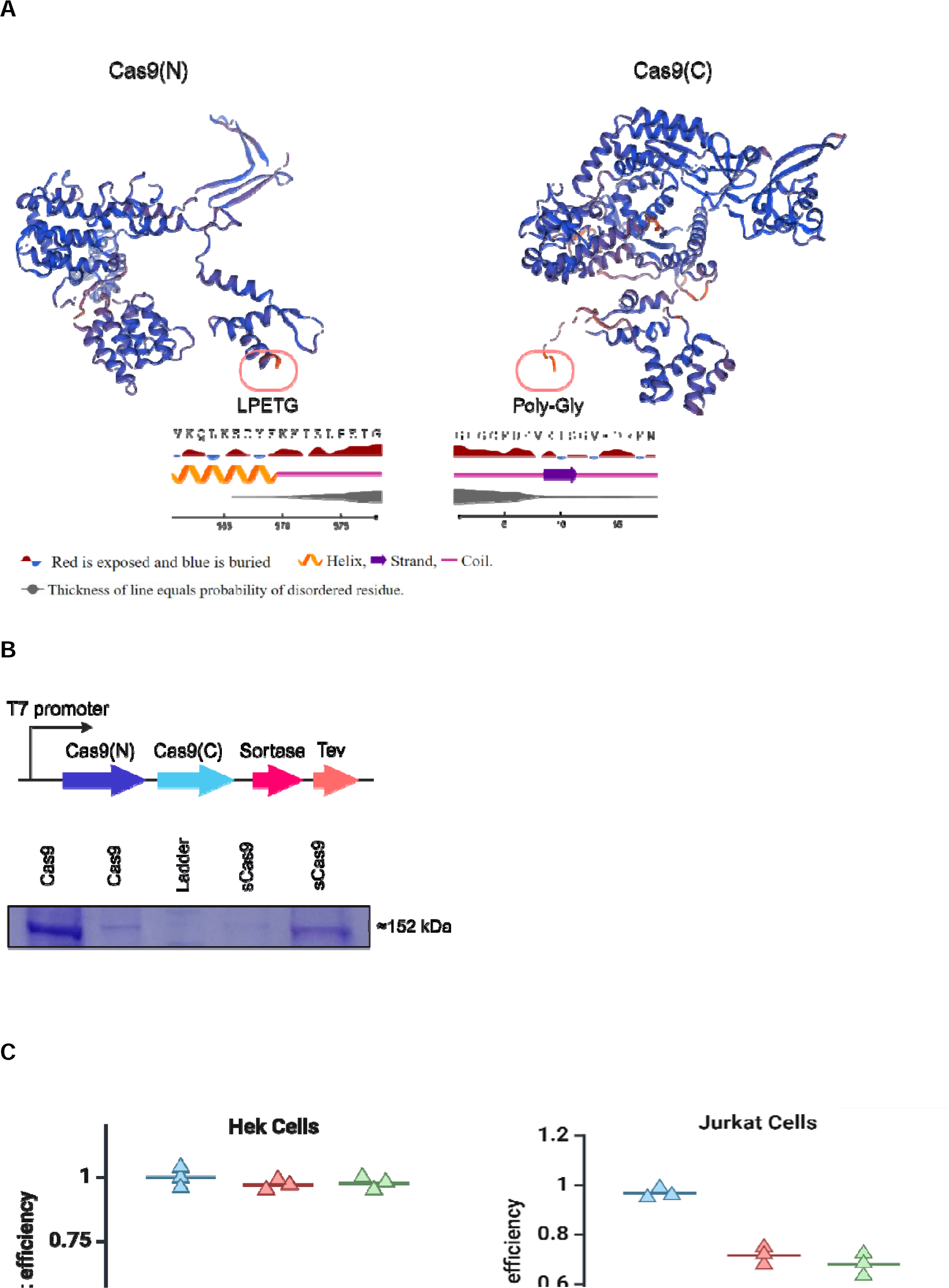
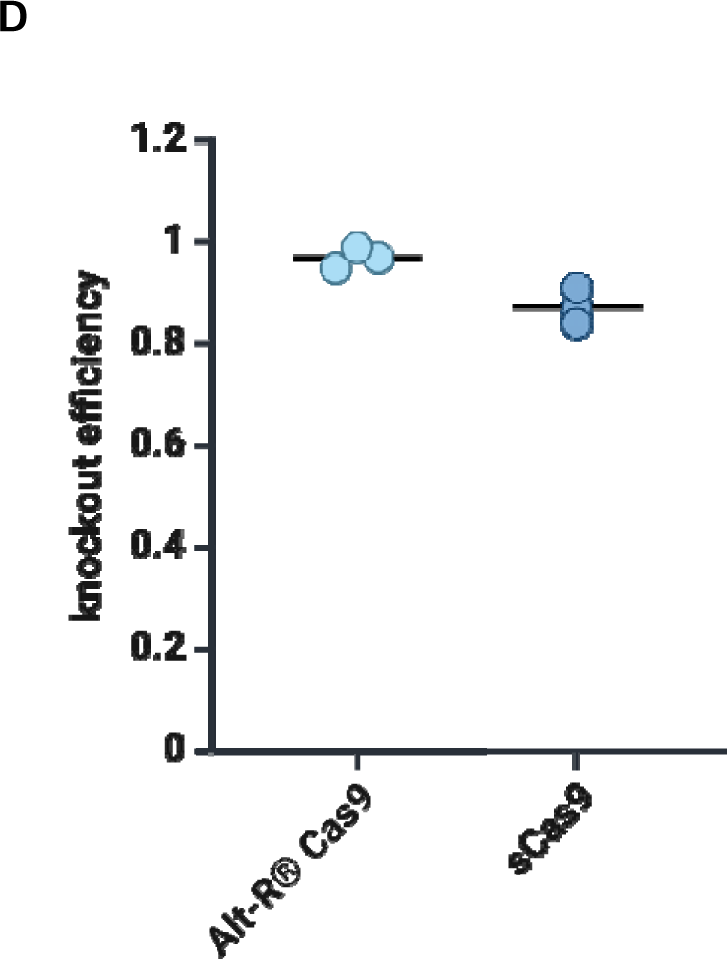
(A) The generated models of the split-Cas9(N) and split-Cas9(C) proteins are depicted in the image. The LPETG and poly-Gly regions are indicated, and the accessibility of these regions was evaluated. (B) The expression cassette map utilized for testing the system in *E. coli*, along with the SDS-PAGE image of the purified Cas9 enzymes from *E. coli*. The Cas9 columns depict the normal Cas9 enzymes expressed in *E. coli*, while the sCas9 proteins represent the enzymatic assembled split-Cas9 enzymes using the sortase-based system in *E. coli*. (C) The knockout efficiency of the designed and synthesized gRNAs was assessed for targeting the *luciferase* gene in the Hek-Luc cell line and the *itgal* gene in Jurkat cells. In the Hek-Luc cell line, the control was normal Hek cells lacking the luciferase gene, while in the Jurkat cells, the control cells were not labeled cells for the Itgal protein. (D) Comparison between the activity of the enzymatically assembled split-Cas9 enzyme (sCas9) produced in *E. coli* and the commercial Alt-R Cas9 enzyme.

Based on our *in silico* findings, we decided to cleave the Cas9 sequence at amino acid position 573, between Glu573 and Cys574. The sortase recognition motif LPETG was added to the C-terminus of the N-terminal Cas9 (Cas9(N)). Additionally, to render the C-terminal Cas9 (Cas9(C)) amenable to sortase-mediated ligation, a Tev protease recognition site (ENLYFQ/G) was introduced at its N-terminus. This motif facilitates the generation of a poly-glycine region in the N-terminus post-cleavage by the Tev protease. These modifications enable sortase-mediated ligation of Cas9(N) and Cas9(C) upon co-expression (Figures 1 and 2A).

### Assessment of the Sortase-mediated assembly system in *E. coli*

To assess the potential of the designed system and confirm its ability to facilitate the ligation of the two split-Cas9 halves into a functional enzyme, we initially conducted experiments in *E. coli*. Employing a one-vector system approach, the two split-Cas9 fragments, along with the assembly machinery (SrtA+Tev), were expressed using the pET-srt split573 plasmid in *E. coli* BL21. The successful expression of the two fragments and subsequent ligation of the full Cas9 enzyme through sortase-mediated ligation was confirmed through Ni-NTA purification and SDS-PAGE. As a control, *E. coli* BL21 was transformed with the pET-srt Cas9 plasmid to express the full Cas9 enzyme. Our SDS-PAGE results demonstrated that the system could successfully ligate the Cas9(N) and Cas9(C) split proteins, with the assembled protein appearing at approximately 152 kDa, corresponding to the size of the Cas9 protein, thus confirming the successful assembly of the split-Cas9 proteins in *E. coli*. Subsequently, we purified this protein using Ni-NTA followed by size exclusion chromatography to enhance its purity. Additionally, we purified the expressed Cas9 using the same method for comparison (Figure 2B).

### Assembled split-Cas9 activity assay

To evaluate the efficacy of the designed gRNAs, we synthesized two gRNAs targeting the integrated *luciferase* gene in Hek-Luc cells and two for recognizing the *itgal* gene in Jurkat cells. Initially, to validate the designed gRNAs, the itgal-gRNAs were transfected alongside Alt-R® Cas9 (IDT) via electroporation into Jurkat cells. Similarly, luc-gRNAs were tested with Alt-R® Cas9 in Hek-Luc cells. Successful knockouts (KOs) of both the *itgal* and *luciferase* genes were confirmed through flow cytometry (FC) and luciferase activity assays, respectively. Based on the obtained results, no significant difference was detected between the gRNAs tested, leading to the selection of luc1 and itg1 gRNAs for subsequent experiments (Figure 2C).

Subsequently, we evaluated the activity of the assembled split-Cas9 protein (sCas9) in these cell lines. We generated ribonucleoprotein complexes (RNPs) with purified sCas9 and commercial Alt-R® Cas9 (IDT) enzyme. These RNPs were transfected into cells, and luciferase activity was assessed after 48 hours. Our findings indicated that sCas9 Alt-R® Cas9 exhibited similar activity, with no significant difference observed. This suggests that the assembly process did not adversely affect the enzyme activity. However, Alt-R® Cas9 demonstrated greater activity than our purified enzymes, which could be due to potential negative effects of our purification method on the enzyme or improvements in the commercial product Alt-R® Cas9 (Figure 2D).

### Establishment of the enzymatic assembly system in mammalian cells

To investigate the potential assembly of sCas9 from its split components using the enzymatic assembly system in mammalian cells, we initially validated the expression of split-Cas9 proteins under the control of the CMV promoter in MC plasmids. This was achieved by adding a GFP gene downstream of Cas9(N) and Cas9(C) using a P2A sequence, resulting in the generation of the MC Cas9(C)-GFP and MC Cas9(N)-GFP plasmids. Additionally, the MC TS-gRNA plasmid was equipped with a mCherry marker. Subsequently, the expression of the plasmids in Hek cells was confirmed through fluorescence microscopy (Figure S2A-C). High viability (> 95%) was observed among the cells individually transfected with these MC plasmids, confirming the absence of plasmid-derived toxicity.

Notably, upon finalizing the constructs, MC plasmids were generated by removing the bacterial backbone sequence and then used for subsequent steps of the study.

We then used a 3-plasmid system utilizing MC Cas9(C), MC Cas9(N), and MC TS-luc at a ratio of 2:2:1 µg to investigate the full system in Hek-Luc cells. After 72 hours of transfection of the plasmids into Hek-Luc cells, we examined the presence of full-size sCas9 in the cells through western blot analysis. The western blot results confirmed the presence of assembled sCas9 in the cells. As a negative control, the lysate of Hek-Luc cells transfected with lipofectamine without any foreign DNA was utilized, and a purified Cas9 protein from *E. coli* was used as a positive control (Figure 3A).

**Figure 3.**
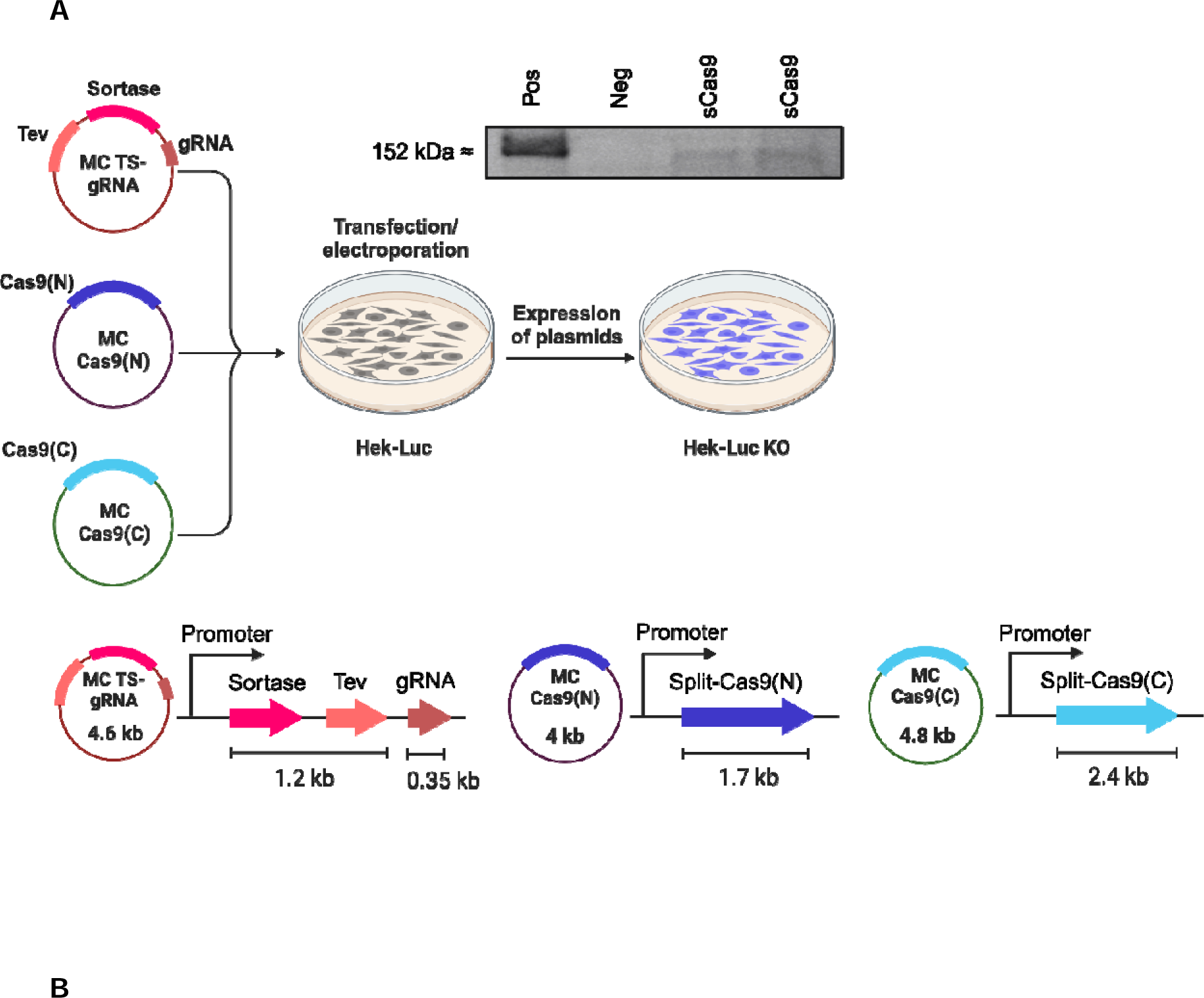
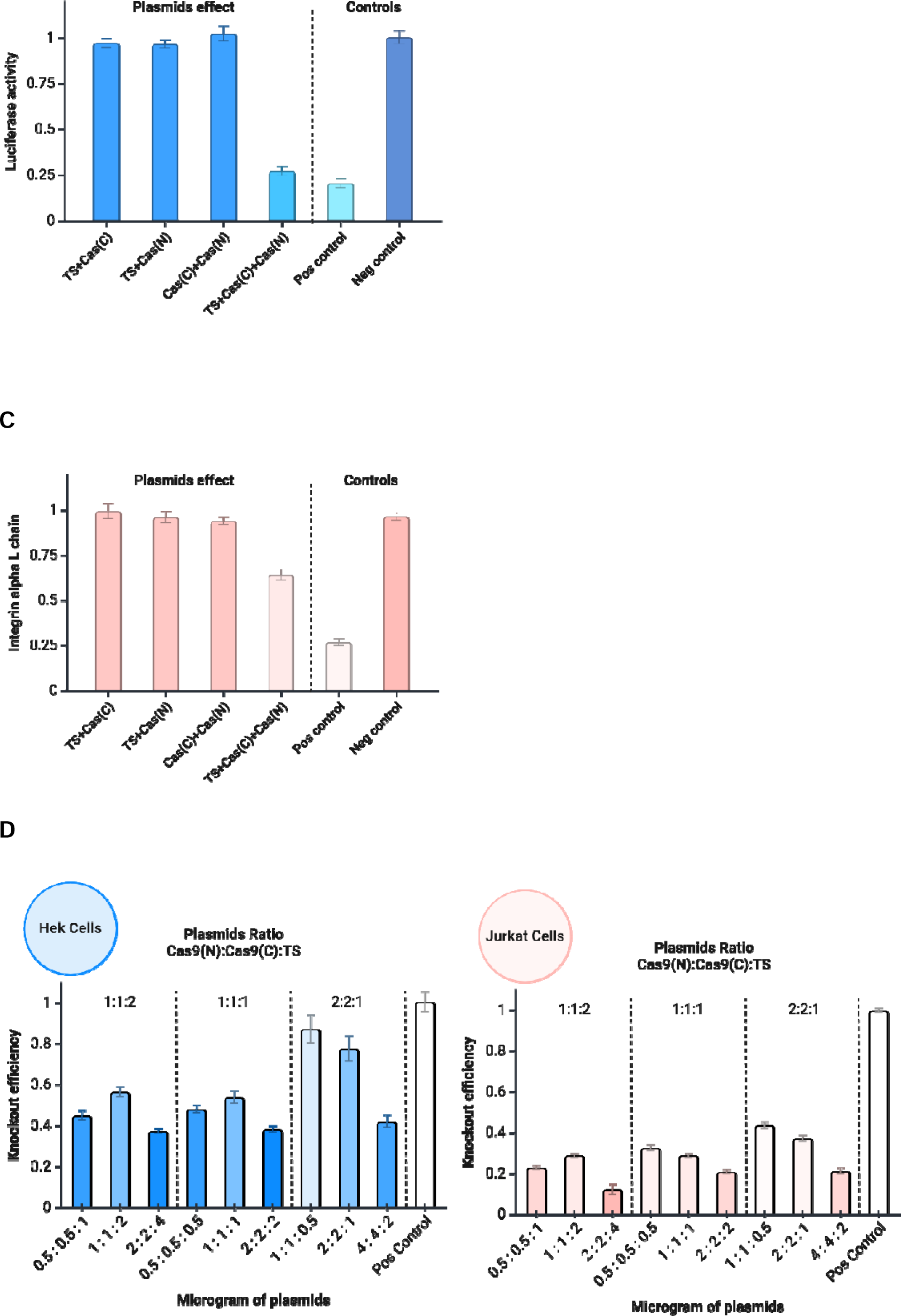
(A) The 3-plasmids system consists of three S/MAR minicircle vectors harboring the split-Cas9 genes, assembly enzymes, and gRNA. The size and map of the cassette in each plasmid are depicted. Additionally, the western blot image shows the enzymatically assembled split-Cas9 enzymes in Hek cells. The positive control was the normal Cas9 enzyme, while the negative control consisted of normal Hek-Luc cells transfected without any plasmid. (B) Luciferase activity in Hek-Luc cells transfected with the constructed plasmids. In this graph, lower activities indicate higher luciferase gene knockout efficiency. Controls included cells transfected with the MC Cas9-luc plasmid carrying a normal Cas9 gene as the positive control and cells transfected with no DNA as the negative control. (C) Labeling of the Itgal protein on the surface of Jurkat cells, which were electroporated with the system’s plasmids to assay the itgal gene knockout efficiency. The number of cells lacking itgal expression was calculated, with the electroporated cells carrying the MC Cas9-itg plasmid used as the positive control. The negative control consisted of electroporated cells without DNA. (D) System optimization to determine the optimal ratio and concentrations of the plasmids for gene knockout in Hek-Luc and Jurkat cells.

Afterwards, we investigated *luciferase* knockout in the transfected Hek-Luc cells with a luciferase activity assay kit. Our results demonstrated a significant reduction in luciferase activity, indicating efficient knockout of the luciferase gene. As a positive control, the MC Cas9-luc plasmid was utilized (Figure 3B).

Then, we further evaluated the functionality of the designed system by targeting the integral membrane protein integrin alpha-L-chain, which is encoded by the *itgal* gene.^28^ Employing the same method, we utilized the 3-plasmid system with split-Cas9 plasmids and MC TS-itg (containing the itg1 gRNA). Similarly, the system effectively knocked out the *itgal* gene, as evidenced by a significant reduction compared to the MC Cas9-itg control plasmid with normal Cas9, which was confirmed through FC analysis (Figure 3C).

### Plasmid concentration and ratio optimization

To study the impact of plasmid ratios in the 3-plasmid system and further optimize the system, we examined different concentrations and ratios of the MC TS-gRNA (assembly) plasmid and split-Cas9 plasmids (MC Cas9(C), MC Cas9(N)). Additionally, we aimed to assess the potential toxicity associated with the transfected plasmids and their effect on cell viability. Plasmids with concentrations ranging from 0.5 µg to 2.5 µg were tested at three different ratios, 1:1:2, 1:1:1, and 2:2:1 for MC Cas9(C) : MC Cas9(N) : MC TS-gRNA plasmids.

The selection of plasmid ratios was based on the hypothesis that cells have a limited capacity for plasmid uptake.^29^ By reducing the amount of the assembly plasmid, there was a higher likelihood that cells would take up all three plasmids necessary for a complete sCas9 enzyme. Furthermore, considering that sortase and Tev can catalyze multiple ligation reactions, it was anticipated that a lower concentration of these enzymes would suffice for a functional split-Cas9 system.

After transfecting Hek-Luc cells with different plasmid ratios and concentrations, we evaluated luciferase activity using the luciferase assay kit after 72 hours. In Jurkat cells, *itgal* knockout was examined by FC analysis 6 days after electroporation (Figure 3D).

Based on the obtained results, it is evident that plasmid ratios characterized by a lower quantity of the assembly plasmid (1:1:0.5 µg and 2:2:1 µg), specifically at a ratio of 2:2:1, generally exhibited superior performance compared to ratios with an equal amount of each plasmid or at a 1:1:2 ratio. This finding underscored the advantage of reducing the amount of the MC TS-gRNA plasmid for effective intracellular uptake of the split-Cas9 halves and subsequent reconstitution of the full sCas9. Moreover, concentrations of 1 µg for split-Cas9 plasmids and 0.5 µg for the assembly plasmid yielded the most favorable outcomes, both in Hek-Luc and Jurkat cells. Furthermore, we assessed cell viability 24 hours post-transfection, and our findings indicated that lower plasmid concentrations did not have any toxic effects during the transfection or electroporation process.

Also, to explore the feasibility of utilizing a 2-plasmid system, we incorporated the *sortase* and *tev* genes along with the gRNA into the MC Cas9(N) plasmid, generating MC Cas9(N)-TS-gRNA. We then evaluated this 2-plasmid system using different concentrations and ratios to optimize its performance and compared it with the 3-plasmid system. Our findings indicated that, in comparison to the 3-plasmid system, the 2-plasmid system demonstrated enhanced potential for genome editing, possibly attributed to its higher transfection efficiency. Moreover, a ratio of 1:1 and a concentration of 1 µg of each plasmid exhibited optimal performance in both Hek-Luc and Jurkat cells (Figure 4A and B).

**Figure 4.**
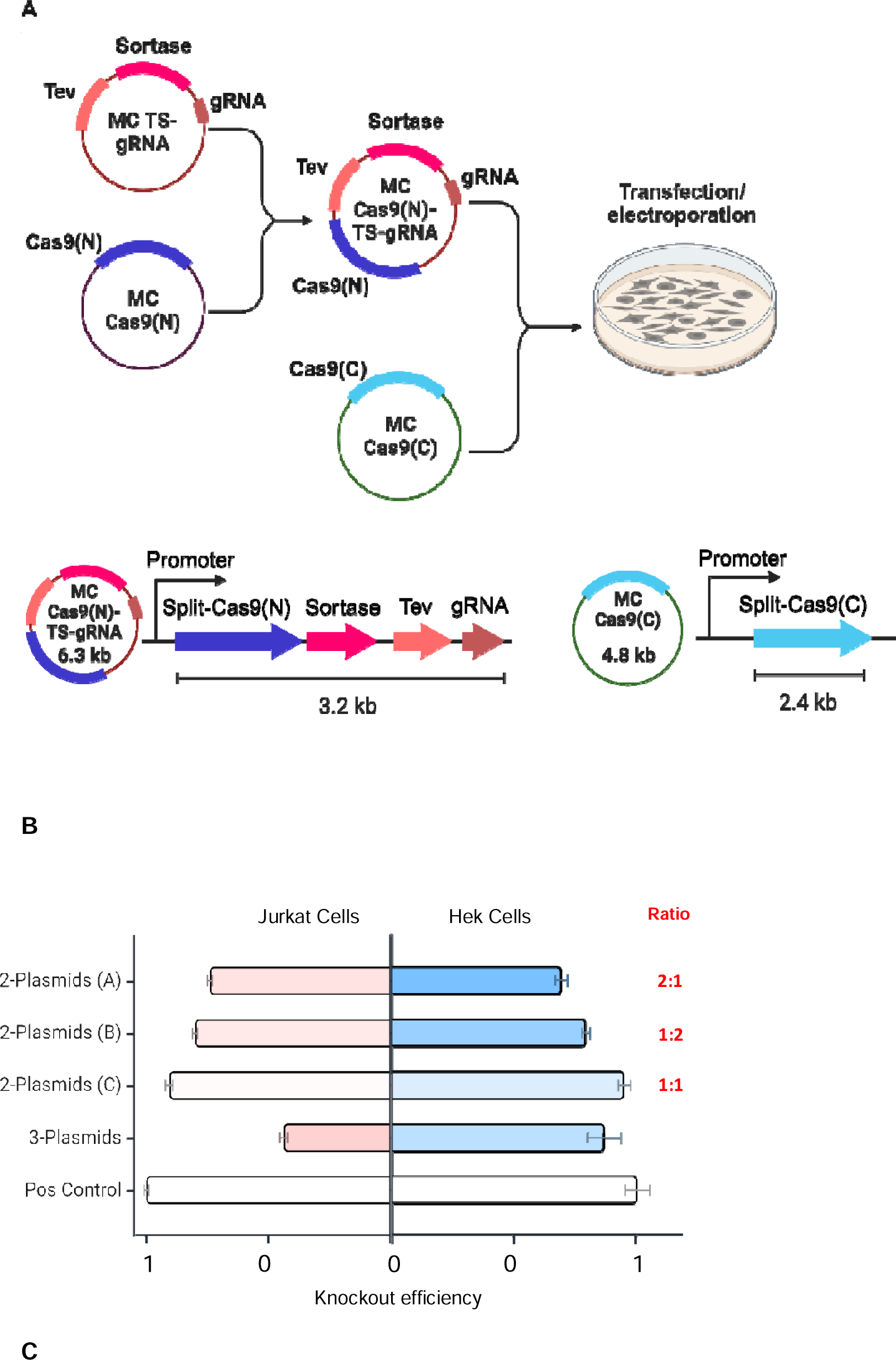
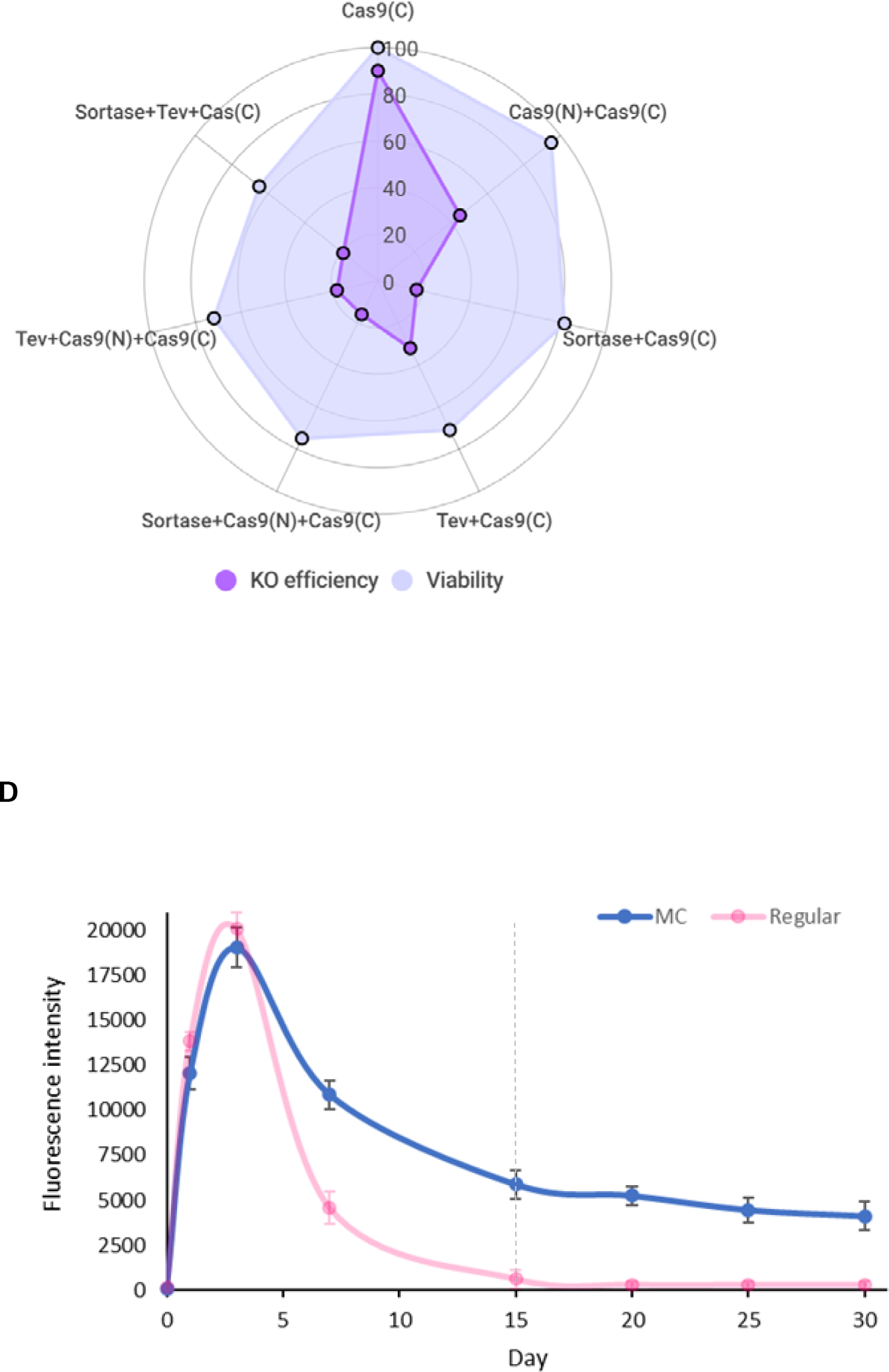
(A) The 2-plasmid system consists of two main plasmids harboring split-Cas9 genes, assembly enzyme-encoding genes, and gRNA. The map, genes, and sizes of each plasmid are depicted. (B) The gene knockout efficiency of the 2-plasmid system for luciferase and itgal genes in Hek-Luc and Jurkat cells, respectively. Additionally, different ratios of the two plasmids, including 2:1, 1:2, and 1:1 µg, were tested. (C) The effect of adding a nuclear localization signal (NLS) to different components of the system on gene knockout efficiency was assessed. Additionally, cell viability was evaluated 24 hours after transfection. The names of the proteins with NLS are labeled on each axis. (D) Stability assay of Hek cells transfected with MC plasmids over a 30-day period with GFP expression. The same plasmid with the bacterial backbone and without the S/MAR region was used for control as a regular plasmids.

### The effect of NLS on the components of the system

The impact of the nuclear localization signal (NLS) on the efficiency of the Cas9 enzyme in genome editing is crucial.^30^ Thus, we aimed to investigate whether performing the assembly process in the nucleus by adding NLS to the assembly enzymes would affect the efficiency of the split-Cas9 system. Also, in our initial system, we incorporated only the NLS in the Cas9(C) split protein; hence, we aimed to evaluate whether adding the NLS to the Cas9(N) split protein could enhance the transfer of sCas9 to the nucleus and thereby improve its efficiency.

To evaluate this possibility, we introduced the NLS to the C-terminus of the Sortase and Tev enzymes and the N-terminus of Cas9(N). Afterwards, we investigated different plasmid combinations in Hek-Luc cells to determine the effect of NLS incorporation in the system and *luciferase* gene knockout. Furthermore, we assessed cell viability 24 hours post-transfection to assess the toxicity of these new plasmids and the impact of delivering higher concentrations of the components to the nucleus.

Based on the luciferase assay results, the various plasmid combinations of the split-Cas9 system demonstrated different knockout efficiencies, ranging from 16% to 90%. The results revealed that our basic system, featuring only an NLS in the C-terminus of the Cas9(C) protein, exhibited the highest knockout efficiency and viability (Figure 4C). The knockout efficacy decreased to less than 50% in the presence of NLS in both the Cas9(C) and Cas9(N) split proteins, possibly due to the rapid transport of the split proteins to the nucleus before assembly but without any significant effect on cell viability.

Furthermore, the addition of NLS to both the SrtA and Tev enzymes decreased both the knockout efficiency and cell viability of the transfected cells. In Hek-Luc cells, this observation suggested that it is more advantageous to assemble the split-Cas9 proteins in the cytosol. Notably, having an NLS in the Cas9(C) protein alone appears to be sufficient for the effective transfer of sCas9 to the nucleus (Figure 4C).

### S/MAR minicircle plasmids

In our study, we opted to employ S/MAR minicircle (MC) plasmids as an alternative to viral vectors for expressing the components of our designed technology. First, we assessed our method for generating MC plasmids, and our experiments confirmed the successful removal of the bacterial backbone sequence. This was confirmed through agarose gel electrophoresis of the purified plasmids after digestion (Figure S3A). Then, we conducted a stability test of the plasmid for GFP expression in Hek cells over 30 days. Next, we compared the MC plasmid to the same plasmid with the bacterial backbone and without S/MAR segments and observed that the presence of the bacterial backbone did not adversely affect the gene expression by our plasmids over a short period of time. However, MC plasmids exhibited stable expression within 30 days, whereas the expression of the regular plasmid dropped after 7 days and disappeared within less than 15 days (Figure 4D).

### Integration of inducible promoters and co-delivery of mRNA and proteins

To evaluate the compatibility of the system with the co-delivery of mRNA and proteins as well as the integration of inducible promoters, we initially expressed and purified the Sortase enzyme (Figure S3B). Subsequently, we encapsulated the Sortase and Tev proteins using lipid-based nanocarriers to mimic protein delivery methods. These nanocarriers, containing the assembly enzymes, were then introduced into Hek-Luc cells that had been transfected with split-Cas9 proteins 48 hours prior, and the luciferase activity was measured 72 hours later. Our findings indicated that the efficiency of the delivery of enzymes via nanocarriers was similar to that of the control, which was the 3-pasmid system. Furthermore, we demonstrated the feasibility of utilizing mRNA encoding the Cas9(N) protein and delivering it to cells expressing the other components of the system. In addition, we studied the combined delivery of mRNA and proteins, which showed an efficiency of approximately 73% compared to that of normal Hek cells without the *luciferase* gene (Figure 5A-C).

**Figure 5.**
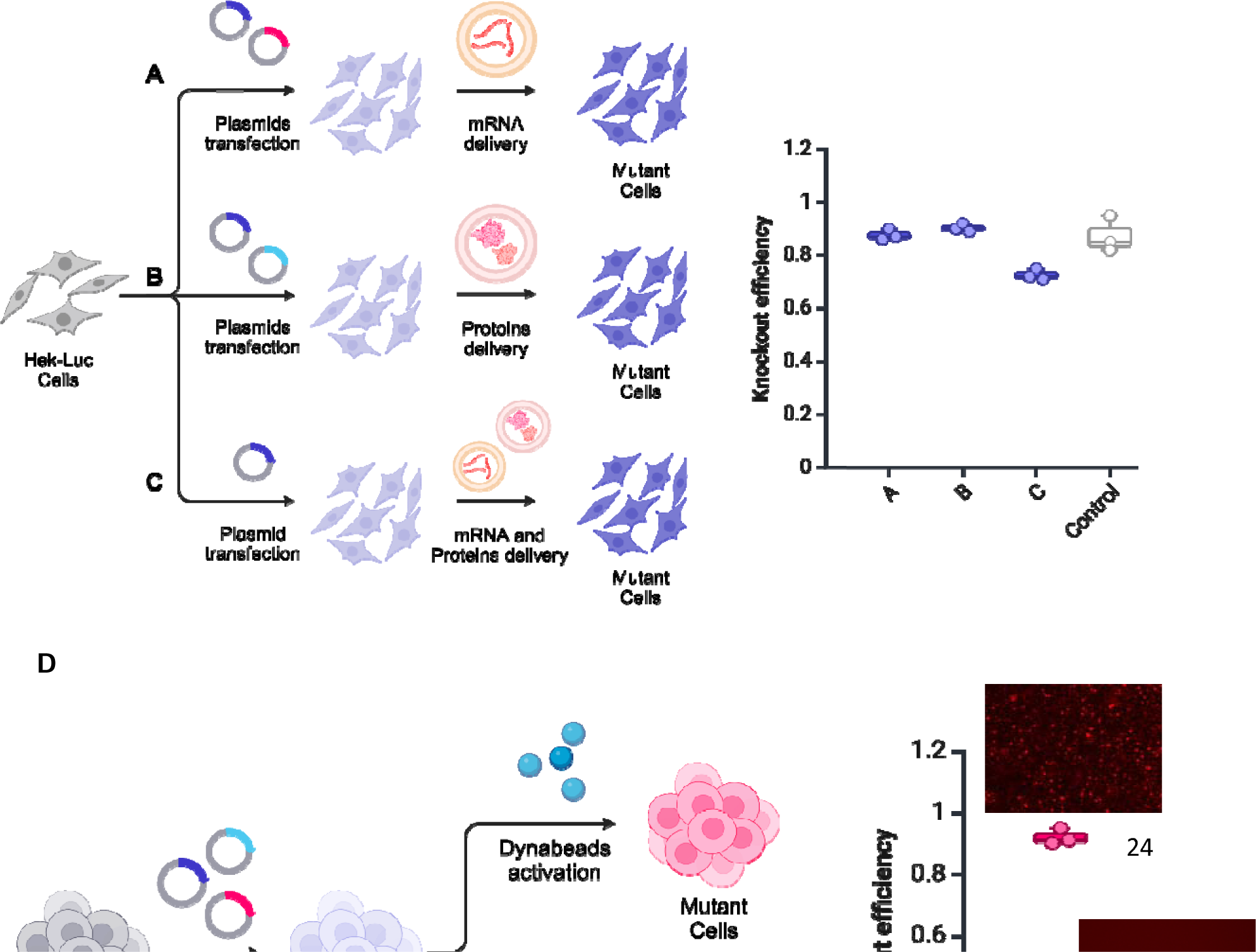
(A-C) Different approaches for delivery of the system components in mRNA and protein forms and assessment of their knockout efficiencies. (A) Cas9(N) mRNA was delivered to cells expressing Cas9(C) and assembly enzymes. (B) The potential of delivering the Sortase and Tev proteins to cells expressing Cas9(N) and Cas9(C). (C) Additionally, the Sortase and Tev proteins were co-delivered with Cas9(N) mRNA to cells containing the Cas9(C) gene. The 3-plasmid system was used as a control, and all the results were compared to the normal Hek cells without the luciferase gene. (D) The CMV promoter upstream of the assembly enzymes and mCherry genes was substituted with the inducible NFAT promoter. The plasmids were electroporated into Jurkat cells, which were then activated using Dynabeads to induce the NFAT promoter. The knockout efficiency was assessed, and the induction of the promoter was confirmed by observing mCherry expression in the cassette using fluorescence microscopy.

Also, to explore the potential of using inducible promoters in this system, we introduced the nuclear factor of activated T cells (NFAT) inducible promoter ^31^ upstream of the sortase, tev, and mCherry genes to generate the MC NFAT-TS-itg plasmid. To induce the NFAT promoter in Jurkat cells, we utilized Dynabeads™ (CD3-CD28). Then, our system’s performance was assessed by co-expressing the MC Cas9(C) and MC Cas9(N) plasmids, each driven by a CMV promoter, along with the MC NFAT-TS-itg assembly plasmid. Given the presence of mCherry in the MC NFAT-TS-itg plasmid construct, we confirmed NFAT activation and mCherry expression using fluorescence microscopy 24 hours post-activation.

Our fluorescence microscopy analysis revealed strong activation of Jurkat cells, leading to the expression of assembly enzymes and mCherry genes under the NFAT promoter. However, no mCherry expression was detected in the uninduced cells. The knockout efficiency in our approach, which utilized the NFAT inducible promoter in activated Jurkat cells, reached approximately 92% compared to the use of the CMV promoter in this system. A comparison between the induced and uninduced NFAT promoters yielded significant results (Figure 5D). These experiments highlight the potential of our system to be precisely controlled by specific promoters for genome editing, demonstrating successful gene knockout.

Notably, with regard to the use of the NFAT promoter, although mCherry expression was not detected in non-activated Jurkat cells, qPCR revealed leakage from the promoter. Nevertheless, the difference observed in the split system under this promoter was significant. These experiments suggest that various promoters can be utilized to tightly control genome editing, including specific promoters for each component of the system.

## Discussion

In this study, we generated a new enzymatic alternative for split-Cas9 technology by utilizing the Sortase A (SrtA) enzyme. Due to the small size of the assembly enzymes, 450 bp for SrtA and 702 bp for Tev, this system can be employed with split-Cas9 proteins in different delivery systems, including adenoviral vectors, to address the issue of the large size of the Cas9 enzyme. Furthermore, the system showed a robust assembly capability with a high efficiency of gene knockout in comparison to regular Cas9 expression in the cells, reaching 98%.

To demonstrate the potential of the assembly system in different cell types, we examined the system in *E. coli*, Hek cells, and Jurkat cells. The high activity of the assembled Cas9 enzymes in all of the abovementioned cells revealed the system’s compatibility with diverse cellular environments, suggesting its applicability in both bacterial and mammalian cells. Additionally, Jurkat cells serve as a reliable model for T lymphocyte cells, and the successful assembly and gene knockout of this system demonstrated its potential for genome editing in T cells. Moreover, the investigated enzymatic system does not require any chemical molecules for split-Cas9 reconstitution. Additionally, having the assembly enzymes in the system enables us to control the assembly and activity of Cas9 in targeted cells, preventing off-target effects.

In this investigation, for selecting the appropriate sites in this system to cleave the Cas9 enzyme, we identified position 573 as the optimal cleavage site through *in silico* analysis. Our *in silico* analysis demonstrated that this position has superior stability and accessibility that is necessary for the ligation of proteins by the SrtA enzyme (Figures 2A and S1). A Sortase recognition motif (LPETG) and a Tev protease recognition site (ENLYFQ/G) in the split-Cas9 proteins enabled successful ligation of the Cas9(N) and Cas9(C) proteins (Figure 1). Our initial results in *E. coli* confirmed the successful assembly of split-Cas9, paving the way for further investigation in mammalian cells (Figure 2B). Subsequently, the system was optimized to determine the best concentrations and ratios of the plasmids for transfection and electroporation into Hek-Luc and Jurkat cells.

The findings revealed that a ratio of 1:1:0.5 µg of Cas9(C):Cas9(N):TS-gRNA plasmids was the ideal ratio and concentration for the 3-plasmid system in both Hek-Luc and Jurkat cells (Figure 3D). Moreover, in the 2-plasmid system, a plasmids concentration of 1:1 µg was the most effective (Figure 4B). Comparing the 3- and 2-plasmid systems revealed higher efficiency with the 2-plasmid system. This phenomenon was possibly attributed to a greater probability of all the system plasmids being transfected into the cells.

Moreover, the effect of adding NLS to the components of the system was tested to assess the potential of performing the assembly process in the nucleus. However, our system decreased the efficiency of gene knockout when we transferred all components, including assembly enzymes and split-Cas9 proteins, into the nucleus. Furthermore, other layouts of NLSs in these components were studied; nevertheless, all of those conditions reduced the knockout efficiency in the Hek-Luc cell line (Figure 4C).

Despite the compact size of the designed system was compatible with adenoviral vectors, this study investigated the use of S/MAR minicircle plasmids for the expression of the genes incorporated into the system. We aimed to address concerns associated with viral vectors, such as limited capacity, potential immune responses, manufacturing complexities, and risk of insertional mutagenesis, by using MC vectors^16,17^. These episomal plasmids facilitated stable gene expression without the use of viral vectors, and we tested gene expression with these plasmids within 30 days. The results indicated that the expression of the designed GFP plasmid was maintained throughout the 30 days, while the expression of the control plasmid declined after 7 days (Figure 4D). MC allows flexible gene expression under a range of conditions and with various promoters, making it a valuable asset for experimental setups requiring controllable gene expression dynamics.

Moreover, our study explored the compatibility of the system for the co-delivery of mRNA and proteins using lipid-based nanocarriers. The delivery of Sortase and Tev proteins via nanocarriers achieved similar efficiency to that of the conventional 3-plasmid system in Hek-Luc cells. Additionally, we demonstrated the feasibility of utilizing mRNA encoding the Cas9(N) protein to enhance the versatility of our system (Figure 5A-C). This feature can be utilized in tightly controlled combined methodologies within medical CRISPR-Cas9 applications as well as in developing various mouse models for studying drug delivery methods or generating engineered model cell lines.

Furthermore, the incorporation of the nuclear factor of activated T cells (NFAT) inducible promoter in Jurkat cells displayed the potential of employing inducible or specific promoters in our system. The significant knockout efficiency, approximately 92% compared to the CMV promoter, highlighted the utility of specific promoters for regulating genome editing via the developed technology (Figure 5D). As a perspective, this capability can be utilized in the genome manipulation of immune cells, particularly in the engineering of chimeric antigen receptor (CAR) T cells under specific conditions. For instance, our system can be developed for application in tumor microenvironments where specific promoters are activated by antigens, hypoxia, low pH, and inhibitors.

Overall, we explored the use of the Sortase A enzyme for protein assembly and post-translational modifications in mammalian cells, particularly in Hek and Jurkat cell lines, and developed an enzymatic-based split-Cas9 system in S/MAR minicircle plasmids. This enzymatic-based approach and our findings present a promising avenue for advancing CRISPR-Cas9 technology toward safer and more effective genome editing applications, as well as for generating new mouse or cell line models as perspectives.

## Acknowledgements

This work was supported by Novo Nordisk Foundation of Denmark, grant NNF21OC0066562.

## Author contributions

Seyed Hossein Helalat: Conceptualization, Methodology, Investigation, Writing-Original draft, Writing – Review & Editing. Helga Thora Kristinsdóttir: Methodology, Investigation, Writing-Original draft. Astrid Dolinger Petersen: Methodology, Investigation, Writing-Original draft. Rodrigo Coronel Téllez: Methodology, Investigation, Writing– Review & Editing. Mads Nordlund Boye: Methodology, Investigation. Yi Sun: Supervision, Writing-Reviewing and Editing, Funding acquisition.

## References

1. Yu, Y., Wu, X., Guan, N., Shao, J., Li, H., Chen, Y., Ping, Y., Li, D., and Ye, H. (2020). Engineering a far-red light-activated split-Cas9 system for remote-controlled genome editing of internal organs and tumors. Sci Adv 6. 10.1126/sciadv.abb1777.

2. Kostyushev, D., Kostyusheva, A., Brezgin, S., Ponomareva, N., Zakirova, N.F., Egorshina, A., Yanvarev, D. V., Bayurova, E., Sudina, A., Goptar, I., et al. (2023). Depleting hepatitis B virus relaxed circular DNA is necessary for resolution of infection by CRISPR-Cas9. Mol Ther Nucleic Acids 31. 10.1016/j.omtn.2023.02.001.

3. Zhao, Z., Shang, P., Mohanraju, P., and Geijsen, N. (2023). Prime editing: advances and therapeutic applications. Trends Biotechnol 41, 1000–1012. 10.1016/j.tibtech.2023.03.004.

4. Liu, Z., Shi, M., Ren, Y., Xu, H., Weng, S., Ning, W., Ge, X., Liu, L., Guo, C., Duo, M., et al. (2023). Recent advances and applications of CRISPR-Cas9 in cancer immunotherapy. Mol Cancer 22, 35. 10.1186/s12943-023-01738-6.

5. Doudna, J.A. (2020). The promise and challenge of therapeutic genome editing. Nature 578, 229–236. 10.1038/s41586-020-1978-5.

6. Zhang, X.-H., Tee, L.Y., Wang, X.-G., Huang, Q.-S., and Yang, S.-H. (2015). Off-target Effects in CRISPR/Cas9-mediated Genome Engineering. Mol Ther Nucleic Acids 4, e264. 10.1038/mtna.2015.37.

7. Truong, D.J.J., Kühner, K., Kühn, R., Werfel, S., Engelhardt, S., Wurst, W., and Ortiz, O. (2015). Development of an intein-mediated split-Cas9 system for gene therapy. Nucleic Acids Res 43. 10.1093/nar/gkv601.

8. Tsuchida, C.A., Brandes, N., Bueno, R., Trinidad, M., Mazumder, T., Yu, B., Hwang, B., Chang, C., Liu, J., Sun, Y., et al. (2023). Mitigation of chromosome loss in clinical CRISPR-Cas9-engineered T cells. Cell 186, 4567–4582.e20. 10.1016/J.CELL.2023.08.041.

9. Kazemian, P., Yu, S.-Y., B. Thomson, S., Birkenshaw, A., R. Leavitt, B., and J. D. Ross, C. (2022). Lipid-Nanoparticle-Based Delivery of CRISPR/Cas9 Genome-Editing Components. Mol Pharm 19, 1669–1686. 10.1021/acs.molpharmaceut.1c00916.

10. Przybyszewska-Podstawka, A., Czapiński, J., Kałafut, J., and Rivero-Müller, A. (2023). Synthetic circuits based on split Cas9 to detect cellular events. Sci Rep 13. 10.1038/s41598-023-41367-z.

11. Pan, Y., Yang, J., Luan, X., Liu, X., Li, X., Yang, J., Huang, T., Sun, L., Wang, Y., Lin, Y., et al. (2019). Near-infrared upconversion-activated CRISPR-Cas9 system: A remote-controlled gene editing platform. Sci Adv 5. 10.1126/sciadv.aav7199.

12. Wright, A. V., Sternberg, S.H., Taylor, D.W., Staahl, B.T., Bardales, J.A., Kornfeld, J.E., and Doudna, J.A. (2015). Rational design of a split-Cas9 enzyme complex. Proc Natl Acad Sci U S A 112. 10.1073/pnas.1501698112.

13. Zetsche, B., Volz, S.E., and Zhang, F. (2015). A split-Cas9 architecture for inducible genome editing and transcription modulation. Nat Biotechnol 33, 139–142. 10.1038/nbt.3149.

14. Weinberg, B.H., Cho, J.H., Agarwal, Y., Pham, N.T.H., Caraballo, L.D., Walkosz, M., Ortega, C., Trexler, M., Tague, N., Law, B., et al. (2019). High-performance chemical- and light-inducible recombinases in mammalian cells and mice. Nat Commun 10. 10.1038/s41467-019-12800-7.

15. Yuan, G., Lu, H., De, K., Hassan, M.M., Liu, Y., Li, Y., Muchero, W., Abraham, P.E., Tuskan, G.A., and Yang, X. (2022). An Intein-Mediated Split-nCas9 System for Base Editing in Plants. ACS Synth Biol 11. 10.1021/acssynbio.1c00507.

16. Leikas, A.J., Ylä-Herttuala, S., and Hartikainen, J.E.K. (2023). Adenoviral Gene Therapy Vectors in Clinical Use—Basic Aspects with a Special Reference to Replication-Competent Adenovirus Formation and Its Impact on Clinical Safety. Int J Mol Sci 24. 10.3390/ijms242216519.

17. Trivedi, P.D., Byrne, B.J., and Corti, M. (2023). Evolving Horizons: Adenovirus Vectors’ Timeless Influence on Cancer, Gene Therapy and Vaccines. Viruses 15. 10.3390/v15122378.

18. Almeida, A.M., Queiroz, J.A., Sousa, F., and Sousa, Â. (2020). Minicircle DNA: The Future for DNA-Based Vectors? Trends Biotechnol 38, 1047–1051. 10.1016/j.tibtech.2020.04.008.

19. Bozza, M., De Roia, A., Correia, M.P., Berger, A., Tuch, A., Schmidt, A., Zörnig, I., Jäger, D., Schmidt, P., and Harbottle, R.P. (2021). A nonviral, nonintegrating DNA nanovector platform for the safe, rapid, and persistent manufacture of recombinant T cells. Sci Adv 7. 10.1126/sciadv.abf1333.

20. Bozza, M., Green, E.W., Espinet, E., De Roia, A., Klein, C., Vogel, V., Offringa, R., Williams, J.A., Sprick, M., and Harbottle, R.P. (2020). Novel Non-integrating DNA Nano-S/MAR Vectors Restore Gene Function in Isogenic Patient-Derived Pancreatic Tumor Models. Mol Ther Methods Clin Dev 17. 10.1016/j.omtm.2020.04.017.

21. Li, J., Zhang, Y., Soubias, O., Khago, D., Chao, F.A., Li, Y., Shaw, K., and Byrd, R.A. (2020). Optimization of sortase A ligation for flexible engineering of complex protein systems. Journal of Biological Chemistry 295. 10.1074/jbc.RA119.012039.

22. Podracky, C.J., An, C., DeSousa, A., Dorr, B.M., Walsh, D.M., and Liu, D.R. (2021). Laboratory evolution of a sortase enzyme that modifies amyloid-β protein. Nat Chem Biol 17. 10.1038/s41589-020-00706-1.

23. Biasini, M., Bienert, S., Waterhouse, A., Arnold, K., Studer, G., Schmidt, T., Kiefer, F., Cassarino, T.G., Bertoni, M., Bordoli, L., et al. (2014). SWISS-MODEL: Modelling protein tertiary and quaternary structure using evolutionary information. Nucleic Acids Res 42. 10.1093/nar/gku340.

24. Zhang, Y. (2008). I-TASSER server for protein 3D structure prediction. BMC Bioinformatics 9. 10.1186/1471-2105-9-40.

25. Delgado, J., Radusky, L.G., Cianferoni, D., and Serrano, L. (2019). FoldX 5.0: Working with RNA, small molecules and a new graphical interface. Bioinformatics 35. 10.1093/bioinformatics/btz184.

26. Mirdita, M., Schütze, K., Moriwaki, Y., Heo, L., Ovchinnikov, S., and Steinegger, M. (2022). ColabFold: making protein folding accessible to all. Nat Methods 19. 10.1038/s41592-022-01488-1.

27. Klausen, M.S., Jespersen, M.C., Nielsen, H., Jensen, K.K., Jurtz, V.I., Sønderby, C.K., Sommer, M.O.A., Winther, O., Nielsen, M., Petersen, B., et al. (2019). NetSurfP-2.0: Improved prediction of protein structural features by integrated deep learning. Proteins: Structure, Function and Bioinformatics 87. 10.1002/prot.25674.

28. Zhang, J., Wang, H., Yuan, C., Wu, J., Xu, J., Chen, S., Zhang, C., and He, Y. (2022). ITGAL as a Prognostic Biomarker Correlated With Immune Infiltrates in Gastric Cancer. Front Cell Dev Biol 10. 10.3389/fcell.2022.808212.

29. Mancinelli, S., Turcato, A., Kisslinger, A., Bongiovanni, A., Zazzu, V., Lanati, A., and Liguori, G.L. (2021). Design of transfections: Implementation of design of experiments for cell transfection fine tuning. Biotechnol Bioeng 118. 10.1002/bit.27918.

30. Shui, S., Wang, S., and Liu, J. (2022). Systematic Investigation of the Effects of Multiple SV40 Nuclear Localization Signal Fusion on the Genome Editing Activity of Purified SpCas9. Bioengineering 9. 10.3390/bioengineering9020083.

31. Jutz, S., Leitner, J., Schmetterer, K., Doel-Perez, I., Majdic, O., Grabmeier-Pfistershammer, K., Paster, W., Huppa, J.B., and Steinberger, P. (2016). Assessment of costimulation and coinhibition in a triple parameter T cell reporter line: Simultaneous measurement of NF-κB, NFAT and AP-1. J Immunol Methods 430. 10.1016/j.jim.2016.01.007.

